# A genome-wide screen in *ex vivo* gallbladders identifies *Listeria monocytogenes* factors required for virulence *in vivo*

**DOI:** 10.1101/2024.08.12.607554

**Authors:** Nicole H. Schwardt, Cortney R. Halsey, Madison E. Sanchez, Michelle L. Reniere

## Abstract

*Listeria monocytogenes* is a Gram-positive pathogen that causes the severe foodborne disease listeriosis. Following oral infection of the host, *L. monocytogenes* disseminates from the gastrointestinal tract to peripheral organs, including the gallbladder, where it replicates to high densities. The gallbladder then becomes the primary bacterial reservoir and source of fecally excreted bacteria. Despite its importance in pathogenesis, little is known about how *L. monocytogenes* survives and replicates in the gallbladder. In this study, we assessed the *L. monocytogenes* genes required for growth and survival in *ex vivo* non-human primate gallbladders using a transposon sequencing approach. The screen identified 43 genes required for replication in the gallbladder, some of which were known to be important for virulence, and others had not been previously studied in the context of infection. We evaluated the roles of 19 genes identified in our screen both *in vitro* and *in vivo*, and demonstrate that most were required for replication in bile *in vitro*, for intracellular infection of murine cells in tissue culture, and for virulence in an oral murine model of listeriosis. Interestingly, strains lacking the mannose phosphoenolpyruvate-dependent phosphotransferase system (PTS) permeases Mpt and Mpo exhibited no defects in intracellular growth or intercellular spread but were significantly attenuated during murine infection. While the roles of PTS systems *in vivo* were not previously appreciated, these results suggest that PTS permeases are necessary for extracellular replication during infection. Overall, this study demonstrates that *L. monocytogenes* genes required for replication in the gallbladder also play broader roles in disease.

## INTRODUCTION

*Listeria monocytogenes* is a Gram-positive bacterium known to survive and replicate in a variety of environments, including soil, sludge, and in mammalian hosts where it is the etiologic agent of the severe foodborne illness listeriosis [1]. Humans are exposed to *L. monocytogenes* multiple times per year as it contaminates ready-to-eat foods due to its frequent occurrence in food processing facilities [2]. While the incidence of disease is relatively low compared to other foodborne illnesses, the case fatality rate of invasive listeriosis is 15-22%, making *L. monocytogenes* one of the most deadly bacterial pathogens [2,3]. In addition, it is estimated that 10-12% of healthy adults exhibit asymptomatic fecal shedding of *L. monocytogenes*, potentially contributing to the spread of this deadly pathogen [1].

After ingestion of *L. monocytogenes*-contaminated food, the bacteria first colonize the lumen of the gastrointestinal (GI) tract where they replicate extracellularly, primarily in the cecum and colon [4]. In susceptible hosts, a small portion of the *L. monocytogenes* in the GI tract invade intestinal epithelial cells, replicate intracellularly, and spread to neighboring enterocytes via actin-mediated motility [5]. *L. monocytogenes* then cross the mucosal barrier and gain access to the lamina propria, Peyer’s patches, and draining mesenteric lymph node (MLN). Bacteria disseminate from the GI tract indirectly to the spleen through the MLN and lymphatics, and directly to the liver via the portal vein [5,6]. Subsequently, 1-5 bacteria migrate from the liver through the hepatic ducts to seed the gallbladder, where they replicate extracellularly in the lumen to very high densities [7]. After a meal, the gallbladder contracts and delivers a bolus of bile and bacteria into the small intestines, reseeding the GI tract [8]. Thus, the gallbladder becomes the main reservoir of *L. monocytogenes* during infection and the source of bacteria excreted in the feces [7]. These observations suggest that replication in the gallbladder is important for infection outcomes and potentially pathogen transmission, and yet little is known about the requirements for *L. monocytogenes* colonization and proliferation in this organ.

The gallbladder is a hollow, sac-like organ in which bile is stored and concentrated. Bile is composed of bile salts, cholesterol, phospholipids, and the heme degradation products biliverdin and bilirubin, which give bile its characteristic color. Bile acts as an emulsifier, aiding in the digestion of lipids in food and exhibiting antimicrobial activity by damaging membranes, nucleic acids, and proteins [9]. Despite this, some bacteria have evolved methods of bile detoxification that render them tolerant to bile and capable of colonizing the gallbladder, including *L. monocytogenes*, *Salmonella enterica*, and *Campylobacter jejuni* [10–12]. Following foodborne infection, *S. enterica* replicates in the gallbladder both extracellularly in biofilms on gallstones and intracellularly within epithelial cells [13]. Additionally, fecal shedding from chronic asymptomatic carriers is important for the pathogenesis and transmission of *S. enterica,* the most famous example of which being Typhoid Mary [11]. In contrast, *C. jejuni,* a pathogen that infects both livestock and humans, replicates extracellularly and localizes to the mucosal folds of the gallbladder, although the role of gallbladder colonization in disease remains unclear [14]. These infection strategies are distinct from that of *L. monocytogenes*, which replicates extracellularly in the lumen of the gallbladder [10].

Animal models of infection have been crucial for understanding disease pathogenesis and gallbladder colonization of bacterial pathogens *in vivo*. Murine infection models are most commonly used, but they pose two major limitations. First, there is a severely restrictive bottleneck in which fewer than five *L. monocytogenes* initially seed the gallbladder following either oral or intravenous inoculation [7]. Second, murine gallbladders are extremely small, containing only 5 - 15 µL of biofluid [15]. These limitations render both genetic and biochemical screening approaches to identify *L. monocytogenes* gallbladder colonization requirements in mouse models impractical. Guinea pig models of oral infection have been used to assess *L. monocytogenes* dissemination from the GI tract to peripheral organs [5], but gallbladder colonization was not assessed in this model. Sheep have been used to study *C. jejuni* infection of gallbladders *in vivo*, but these studies are low-throughput and require veterinary surgical expertise to complete [12,14]. Purified bile salts and reconstituted powdered bile are frequently used to mimic the gallbladder environment *in vitro*, but it is not clear what concentrations and diluent accurately represent gallbladder biofluid. In fact, studies investigating *L. monocytogenes* requirements for survival in reconstituted porcine bile and *ex vivo* porcine bile identified distinct genes [16,17], supporting the notion that reconstituted bile does not mimic the biofluid encountered *in vivo*.

In this study, we sought to identify *L. monocytogenes* genes required for replication in the mammalian gallbladder using a transposon sequencing (Tn-seq) approach. This technique combines saturating transposon mutagenesis with next-generation sequencing to assess the contribution of every genetic locus in a high-throughput manner [18,19]. Tn-seq has been used to identify essential genes and genes conditionally essential for survival in a host for several pathogenic bacteria, including *Staphylococcus aureus* [20]*, Vibrio cholerae* [21]*, Streptococcus pneumoniae* [19], and recently *L. monocytogenes* [22]. However, the restrictive bottlenecks in mouse models of listeriosis make the use of global genetic approaches to study gallbladder colonization *in vivo* unfeasible. Here, we performed Tn-seq on *L. monocytogenes* in *ex vivo* non-human primate gallbladders and identified dozens of genes necessary for survival and replication in this environment and more broadly in the context of a murine model of listeriosis.

## RESULTS

### An unbiased approach identifies *L. monocytogenes* genes required for growth and survival in the gallbladder lumen

To identify *Listeria monocytogenes* genes required for gallbladder colonization, we established a novel model of gallbladder colonization using non-human primate (NHP) gallbladders obtained from the Washington National Primate Research Center Tissue Distribution Program. There are several advantages to NHP organs over conventional murine models. First, NHP gallbladders can be inoculated with bacteria via syringe, eliminating the bottlenecks to colonization encountered during murine infections. Second, NHP organs are larger and contain ∼1,000-fold more biofluid than murine gallbladders, which can support more bacterial biomass or be harvested for *in vitro* assays. Finally, organs were obtained from NHPs at the endpoint of other non-infectious experiments, and therefore no additional animals were sacrificed for this model.

In the development of the gallbladder colonization model, we first assessed whether *ex vivo* gallbladder biofluid (bile) supports growth of *L. monocytogenes.* Bile harvested from three independent NHP gallbladders was determined to be sterile and supported exponential growth of *L. monocytogenes in vitro*. We next injected mid-log *L. monocytogenes* into the lumen of intact *ex vivo* gallbladders and monitored bacterial survival over time by removing luminal contents with a syringe and plating to enumerate colony forming units (CFU). Interestingly, we observed consistent reductions in CFU shortly after inoculation, followed by exponential growth that plateaued between 6 and 12 hours post-injection (data not shown). To minimize host cell death and maintain the integrity of the organs, we limited the incubation time to 6 hours in subsequent experiments.

After establishing growth conditions for *L. monocytogenes* in NHP gallbladders, we used this *ex vivo* organ model to investigate the *L. monocytogenes* genes required for survival and growth in the gallbladder lumen using the unbiased global genetic approach of transposon sequencing (Tn-seq). Four NHP gallbladders were inoculated via syringe with a saturated transposon mutant library of *L. monocytogenes*. To monitor growth of the mutant library in the organs, samples of luminal contents were collected 30 minutes and 6 hours post-injection. As observed previously with wild type (WT) *L. monocytogenes*, an initial reduction in CFU at 30 minutes post-injection was followed by exponential growth through 6 hours, with an average doubling time of 46 minutes (Fig 1A). This doubling time is similar to that observed for *L. monocytogenes* growing in rich medium, demonstrating robust growth in this environment. After 6 hours of incubation in the gallbladders, the entire luminal contents were harvested, diluted in brain heart infusion (BHI) broth, and incubated for 2 hours to increase biomass. Bacterial genomic DNA was then isolated and libraries were prepared for Illumina sequencing of the transposon insertion sites (Fig 1B and S1 Table). Using the parameters of a log_2_ fold-change less than - 1.50 and an adjusted *p*-value of less than 0.05, mutants in 43 genes were significantly depleted after incubation in the gallbladders compared to the input libraries, indicating that these genes are required for growth or survival in the NHP gallbladder lumen.

**Figure 1.**
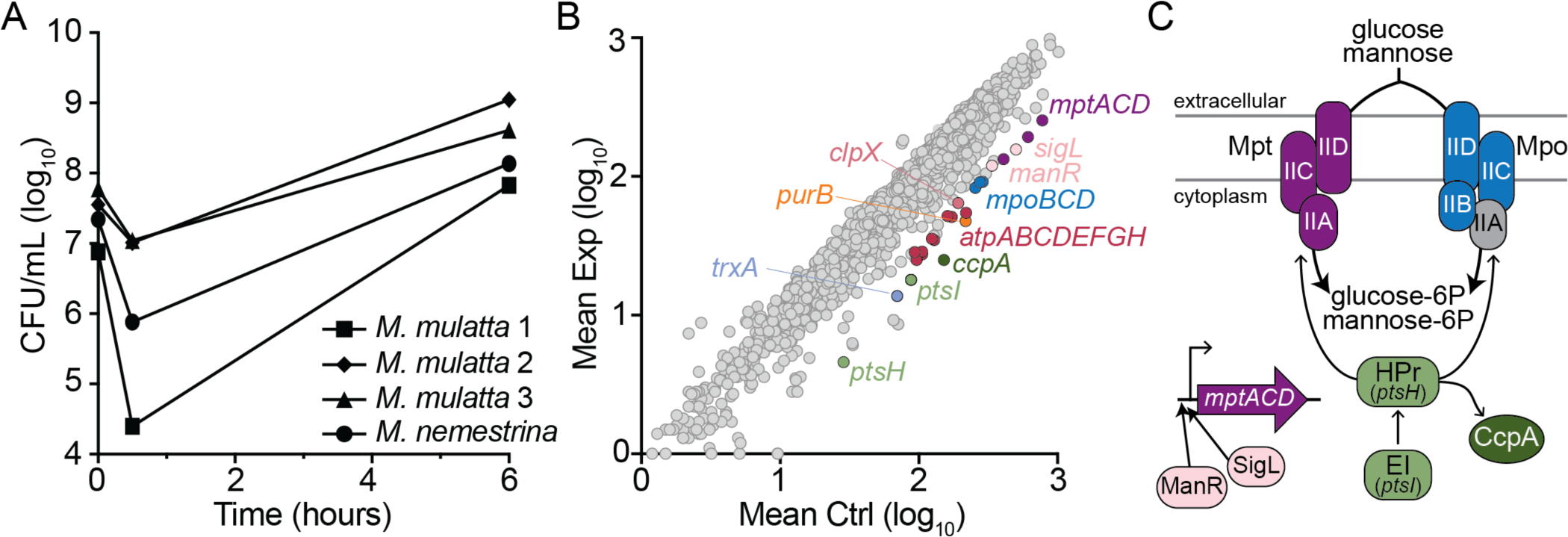
Tn-seq in *ex vivo* NHP gallbladders identifies *L. monocytogenes* genes necessary for growth and survival. (A) Growth of the *L. monocytogenes* transposon library in NHP gallbladders over 6 hours. Gallbladders were inoculated via syringe and bacteria were collected for CFU enumeration 30 minutes and 6 hours post-infection. Gallbladders from three *Macaca mulatta* and one *M. nemestrina* were used. (B) Average read counts of *L. monocytogenes* genes in the gallbladder experimental condition (Exp) compared to the input library (Ctrl). (C) Model depicting the PTSs and their regulators identified by Tn-seq, color-coded to match panel B. Only Mpo-IIA (gray) was not significantly depleted after incubation in NHP gallbladders.

Genes significantly depleted after incubation in the gallbladder included those involved in stress responses, such as *trxA* encoding thioredoxin and *clpX* encoding a Clp protease ATPase subunit, consistent with the known antimicrobial effects of bile. We also identified the adenylosuccinate lyase encoded by *purB*, consistent with the importance of purine biosynthesis in surviving bile stress [17]. Depleted genes also included those encoding two phosphoenolpyruvate-dependent phosphotransferase system (PTS) permeases known to import both glucose and mannose (*mptACD* and *mpoBCD*) [23]. PTSs are multi-protein complexes utilized by many bacteria to import and phosphorylate defined carbohydrates, with the sugar specificity determined by the Enzyme II (EII) complex proteins [24]. While the *L. monocytogenes* genome encodes 29 complete PTSs [23], our screen identified Mpt and Mpo as the only EIIs required for growth in the gallbladder (Fig 1C). In addition to the PTS permeases, genes encoding regulators that activate transcription of the *mpt* and *mpo* operons (*manR* and *sigL*), and for phosphorylation of the PTS sugars (*ptsI* and *ptsH,* encoding EI and HPr, respectively) were also significantly depleted in the gallbladder condition. Identification of multiple PTS-related operons and their regulators, which lie at distinct genetic loci, suggests that *L. monocytogenes* imports glucose and/or mannose via Mpt and Mpo for growth in the gallbladder (Fig 1C). The Tn-seq also identified 8 of the 9 genes encoding the F-type ATP synthase as depleted after growth in the gallbladders. The F-type ATP synthase, while not essential for aerobic growth, is required for anaerobic replication in *L. monocytogenes* [25], suggesting that the *ex vivo* gallbladder may be an oxygen-limited environment. Overall, analysis of our screen identified multiple whole operons as depleted after growth in the NHP gallbladder, indicating that the screen was robust. Importantly, many genes identified here have not been previously implicated in the context of infection.

### Genes identified by Tn-seq in the NHP gallbladder contribute to replication in bile *in vitro*

To investigate the roles of the identified genes in *L. monocytogenes* physiology and pathogenesis, we used allelic exchange techniques to generate nine deletion mutants, representative of 19 genes identified in our screen (Table 1). To evaluate the PTS permeases, the entire *mptACD* (Δ*mpt*) or *mpoABCD* (Δ*mpo*) operons were deleted and a double mutant lacking both operons (*ΔmptΔmpo*) was generated. To evaluate the F-type ATP synthase, the *atpB* open reading frame was deleted, which eliminates functionality of the entire complex [25]. The mutants were then grown individually either in BHI broth or NHP bile in 96-well plates, and CFU were enumerated after 0.5 and 6 hours of static incubation. Because the oxygen status of the gallbladder lumen is not known, *L. monocytogenes* growth was evaluated in both aerobic and anaerobic conditions. Several mutants exhibited general growth defects and replicated significantly less than WT in rich medium, including Δ*ccpA*, Δ*ptsI*, and Δ*trxA* (Fig 2). The Δ*atpB* strain had the most striking phenotype in BHI, displaying a slight ∼4-fold reduction in CFU in the presence of oxygen and a complete lack of growth in anaerobic conditions (Fig 2B). This is consistent with a previous report documenting that the F-type ATPase is essential for *L. monocytogenes* anaerobic growth [25]. When incubated in bile under anaerobic conditions, all mutants were significantly impaired, with the exception of Δ*ccpA* (Fig 2B). This decrease in CFU at 6 hours was not driven by differences in killing early after inoculation, as all strains exhibited similar reductions in CFU at 30 minutes (S1 Fig). Interestingly, some mutants exhibited an oxygen-dependent phenotype. For example, Δ*atpB*, Δ*mpt*, and Δ*mpt*Δ*mpo* grew similarly to WT in bile under aerobic conditions (Fig 2A), but displayed reduced growth under anaerobic conditions (Fig 2B). Together, these results demonstrated that the genes identified by Tn-seq as important during growth in the NHP gallbladder lumen are also required for growth in bile *in vitro*, even in the absence of competing strains.

**Figure 2.**
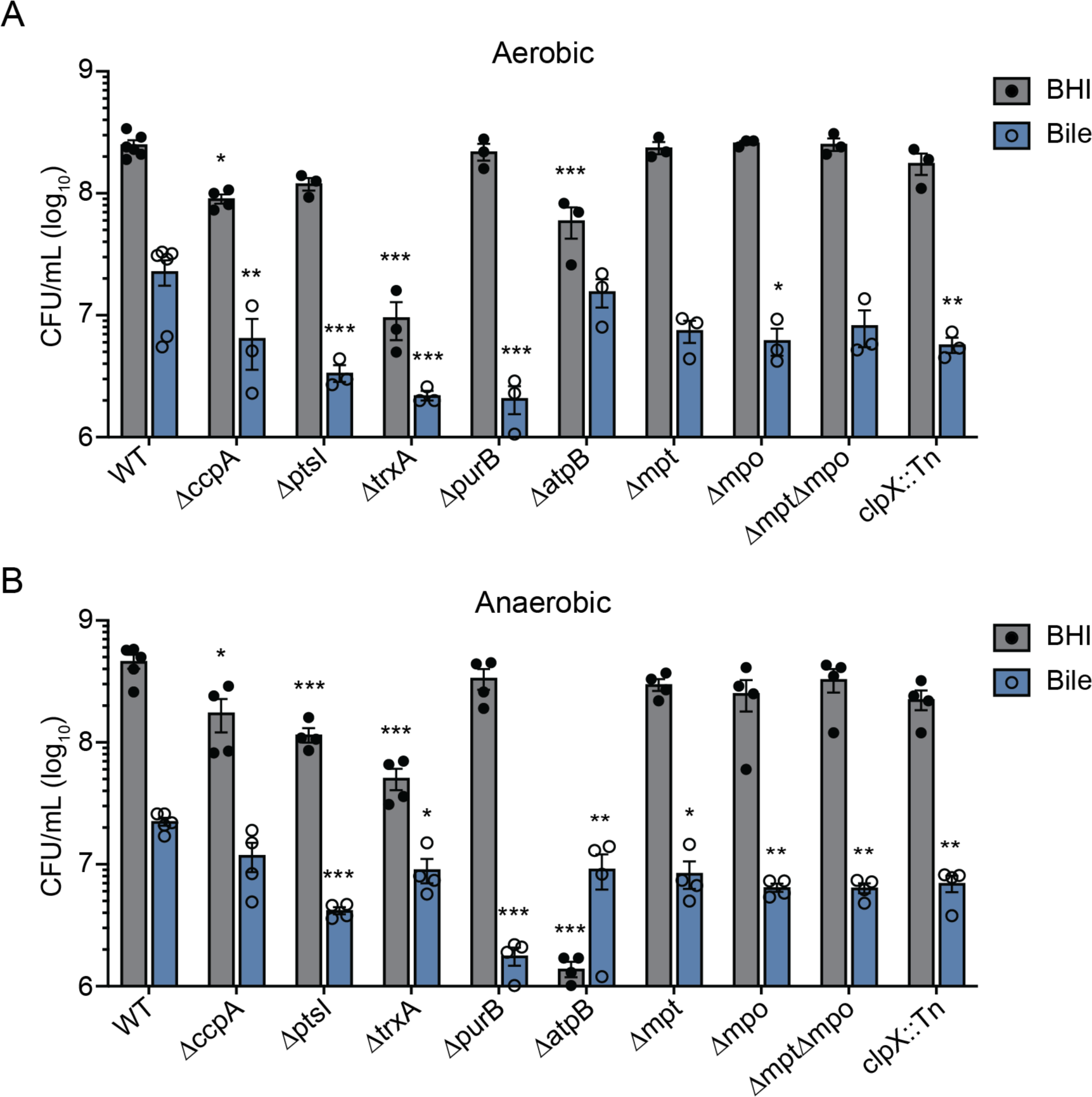
Growth of mutants *in vitro*. BHI or NHP bile was inoculated with 10^6^ CFU/mL of each *L. monocytogenes* strain, incubated for 6 hours statically in an aerobic incubator (A) or in an anaerobic chamber (B), and then CFU were enumerated. Each circle is an individual data point, while the bars indicate the means and the error bars represent the standard error of the mean (SEM) of at least 3 biological replicates. * *p* < 0.05, ** p < 0.01, *** *p* < 0.001, as determined by Dunnett’s multiple comparisons test compared to WT in each medium condition.

**Table 1.** Genes-of-interest depleted after *L. monocytogenes* incubation in NHP gallbladders.

| LMRG | Lmo | Gene Name | Description | log2FC |
| --- | --- | --- | --- | --- |
| LMRG_01368 | lmo1599 | *ccpA* | Catabolite control protein A | -2.56 |
| LMRG_02102 | lmo1002 | *ptsH* | PTS, phosphocarrier protein HPr | -2.42 |
| LMRG_02103 | lmo1003 | *ptsI* | PTS enzyme I (EI) | -2.23 |
| LMRG_00679 | lmo1233 | *trxA* | Thioredoxin | -2.27 |
| LMRG_02498 | lmo1773 | *purB* | Adenylosuccinate lyase | -2.16 |
| LMRG_01718 | lmo2530 | *atpG* | ATP synthase subunit gamma | -1.98 |
| LMRG_01714 | lmo2534 | *atpE* | ATP synthase subunit c | -1.91 |
| LMRG_01719 | lmo2529 | *atpD* | ATP synthase subunit beta | -1.86 |
| LMRG_01720 | lmo2528 | *atpC* | ATP synthase subunit epsilon | -1.85 |
| LMRG_01715 | lmo2533 | *atpF* | ATP synthase subunit b | -1.84 |
| LMRG_01717 | lmo2531 | *atpA* | ATP synthase subunit alpha | -1.78 |
| LMRG_01716 | lmo2532 | *atpH* | ATP synthase subunit delta | -1.74 |
| LMRG_01713 | lmo2535 | *atpB* | ATP synthase subunit a | -1.62 |
| LMRG_02346 | lmo0097 | *mptC/manM* | PTS, mannose-specific IIC component | -1.66 |
| LMRG_02345 | lmo0096 | *mptA/manL* | PTS, mannose-specific IIA/IIB component | -1.61 |
| LMRG_02347 | lmo0098 | *mptD/manN* | PTS, mannose-specific IID component | -1.61 |
| LMRG_00469 | lmo0781 | *mpoD/levG* | PTS, mannose-specific IID component | -1.65 |
| LMRG_00470 | lmo0782 | *mpoC/levF* | PTS, mannose-specific IIC component | -1.61 |
| LMRG_02869 | lmo0783 | *mpoB* | PTS, mannose-specific IIB component | -1.60 |
| LMRG_00718 | lmo1268 | *clpX* | ATP-dependent protease, ATP-binding subunit | -1.57 |

Highlighted genes are predicted to be encoded in an operon [26].

### Genes identified by Tn-seq as critical for survival in the gallbladder lumen also contribute to intracellular fitness

Tn-seq identified *L. monocytogenes* genes required for growth in the gallbladder lumen, which represents one of the extracellular environments that *L. monocytogenes* encounters during infection. While extracellular niches of infection remain largely uncharacterized, the determinants of intracellular infection and their roles in systemic disease have been thoroughly described. The intracellular lifecycle begins with *L. monocytogenes* entering host cells via phagocytosis or receptor-mediated endocytosis, followed by cytosolic replication and intercellular spread to neighboring cells via actin-dependent motility [27]. To determine if the genes we identified also have important roles in the intracellular lifecycle, we assessed cell-to-cell spread and cytosolic replication of each of the *L. monocytogenes* mutants in cell culture.

The intracellular lifecycle was first evaluated via plaque assay in which a monolayer of cells is infected with *L. monocytogenes* and then immobilized in agarose containing gentamicin to prevent extracellular growth. Three days post-infection, the live cells are stained and the zones of clearance formed by *L. monocytogenes* are measured as an indicator of intracellular growth and intercellular spread. In this assay, most mutants formed significantly smaller plaques than those formed by WT, while mutants lacking the PTS operons *mpt* and *mpo* formed plaques similar in size to WT. (Fig 3A). Notably, infections with *ΔatpB* resulted in no visible plaque formation. We hypothesize this is due to the Δ*atpB* requirement for oxygen, which may be limiting in cells with the agarose overlay. Plaque areas were also measured after infection with the complemented strains, in which each deleted gene was expressed from its native promoter at an ectopic site on the chromosome. With one exception, complementation restored plaque areas to WT levels (Fig 3A). The *ptsI* complemented strain produced even smaller plaque areas than Δ*ptsI,* although this strain did restore other Δ*ptsI* growth defects, as discussed below.

**Figure 3.**
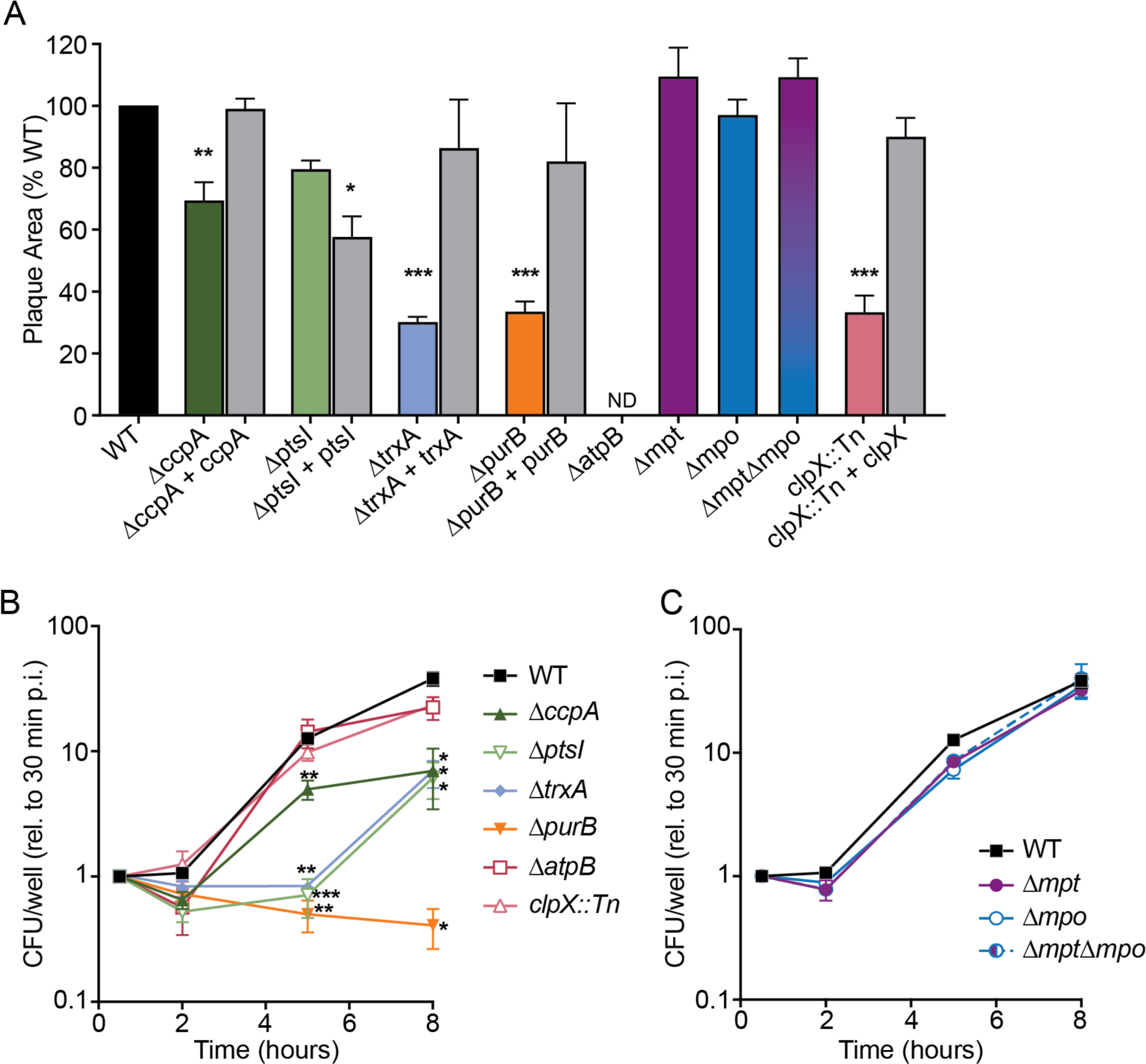
Intracellular replication and intercellular spread. (A) Plaque areas formed in L2 fibroblasts, measured as a percentage of that formed by WT *L. monocytogenes*. * *p* < 0.05, ** p < 0.01, *** *p* < 0.001, as determined by one-way ANOVA with multiple comparisons to the mean of WT. ND, not detected. (B-C) Intracellular growth of mutants in BMDMs, normalized to CFU at 30 minutes for each strain. * *p* < 0.05, ** p < 0.01, *** *p* < 0.001, as determined by Dunnett’s multiple comparisons test compared to WT in each condition. Data in all panels are the means and SEMs of at least 3 independent experiments.

The plaque assay measures both cytosolic replication and cell-to-cell spread. To identify the role of each gene in intracellular growth, we measured replication kinetics in primary bone marrow-derived macrophages (BMDMs) over 8 hours. The *ΔpurB* mutant exhibited the most dramatic phenotype as it did not replicate in the host cytosol (Fig 3B). Several additional mutants displayed attenuated intracellular growth, including *ΔccpA, ΔptsI*, and *ΔtrxA* (Fig 3B). Intracellular growth was fully restored in the complemented strains, including Δ*ptsI* + *ptsI* (S2 Fig). Despite defects in plaque formation, both *clpX::Tn* and *ΔatpB* grew similarly to WT in BMDMs, indicating that these mutants are defective specifically in the cell-to-cell spread stage of the intracellular lifecycle. Finally, strains lacking the PTS operons *(Δmpt, Δmpo, ΔmptΔmpo*) displayed no defects in cytosolic growth, consistent with these strains forming WT-sized plaques (Fig 3C). Together, these results indicated that although we identified these genes using the selective pressure of extracellular growth in a mammalian organ, many also contribute to intracellular infection. However, the PTS operons were found to be dispensable during intracellular infection, consistent with prior work [23].

### *L. monocytogenes* genes important in the NHP gallbladder are required for oral infection of mice

The Tn-seq screen identified many genes important for extracellular replication in an NHP gallbladder as well as intracellular growth and intercellular spread in murine cells. Thus, we hypothesized that these genes would be important for virulence in a mouse model of oral listeriosis. For these infections, 6-7 week old female BALB/c mice were given streptomycin in their drinking water for 2 days and fasted for 16 hours prior to infection to increase susceptibility to oral infection [4,28]. Mice were then fed 10^8^ CFU of each *L. monocytogenes* strain via pipette. Body weights were recorded daily as a measurement of global disease severity. Mice infected with WT lost nearly 20% of their initial body weight throughout the 4 day infection, whereas mice infected with most of the mutant strains exhibited significantly less weight loss (Figs 4A and 4B). Notably, mice infected with Δ*ccpA*, Δ*mpo*, or Δ*mpt*Δ*mpo* lost approximately the same amount of weight as mice infected with WT, suggesting that that these genes may not be required for *L. monocytogenes* pathogenesis *in vivo*. Conversely, mice infected with *ΔptsI*, *ΔtrxA*, and *ΔpurB* lost very little weight over the 4 day infection, suggesting that these strains were severely attenuated in their pathogenicity.

**Figure 4.**
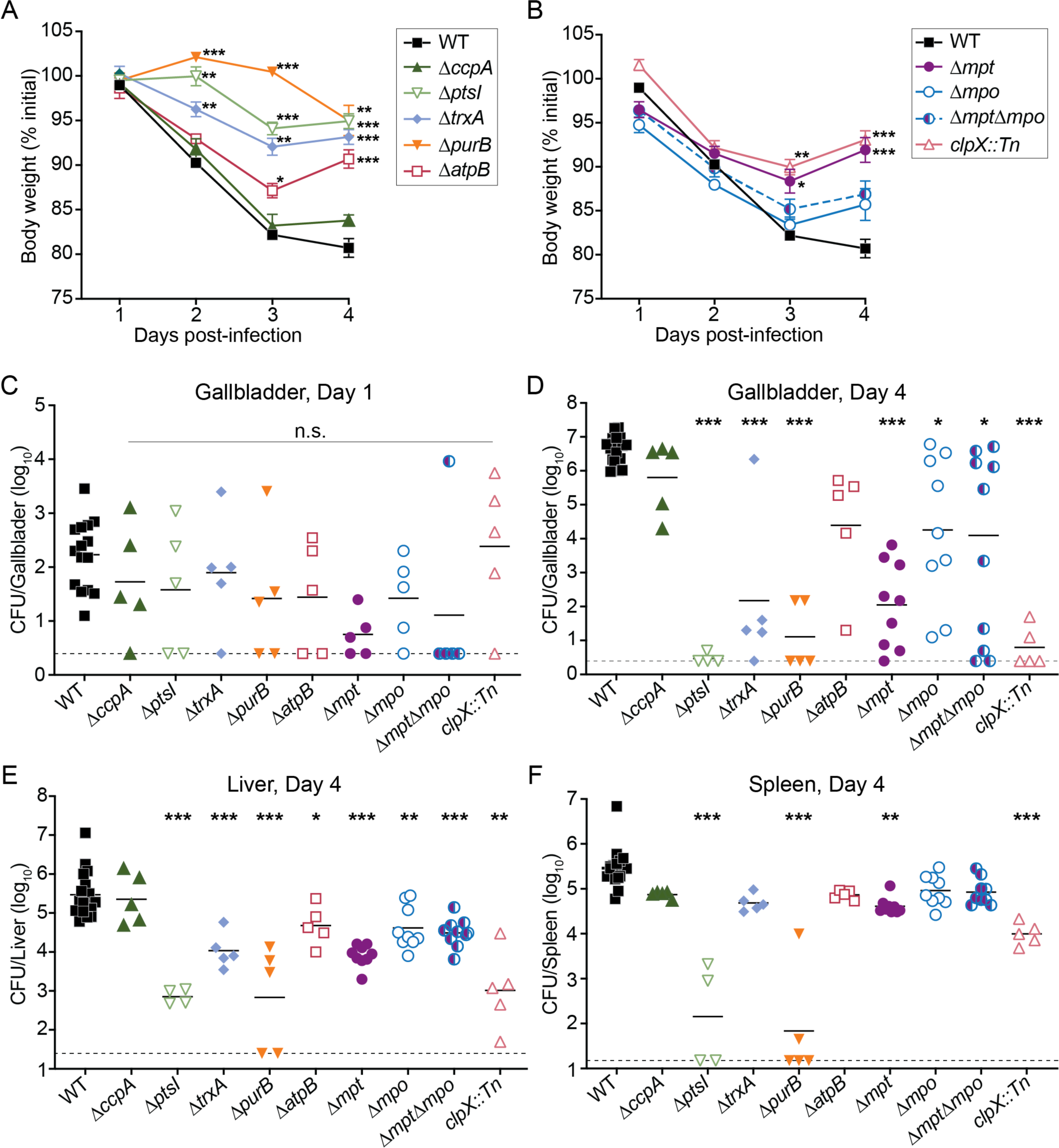
Murine model of oral listeriosis. Mice were orally infected with 10^8^ of each *L. monocytogenes* strain. (A-B) Body weights of infected mice over time, reported as a percentage of initial weight before streptomycin treatment. Data are means and SEM of n=4-33. (C-F) CFU were enumerated from tissues at 1 or 4 days post-infection. Each data point represents a single mouse (n=15-18 for WT, n=4-10 for mutants), horizontal solid lines represent geometric means, and the dotted lines represent the limit of detection. The data are combined from 4 independent experiments. * *p* < 0.05, ** *p* < 0.01, *** *p* < 0.001, as determined by Dunnett’s multiple comparisons test compared to WT in each condition.

To assess bacterial burdens throughout infection, mice were euthanized and CFU were enumerated from organs at both 1 and 4 days post-infection (dpi). At 1 dpi, bacterial burdens in the gallbladders were similar between WT and all mutant strains, indicating that these genes are not required for dissemination from the GI tract to the gallbladder (Fig 4C). By 4 dpi, 7 of the 9 mutants displayed significantly decreased bacterial burdens in the gallbladder compared to WT (Fig 4D). Bacterial burdens in mice infected with *ΔptsI*, *ΔpurB*, and *clpX::Tn* were decreased by more than 300,000-fold compared to mice infected with WT. In contrast, bacterial loads of mice infected with Δ*trxA*, *Δmpt*, *Δmpo*, *ΔmptΔmpo*, and *ΔatpB* displayed more variability in CFU between animals and were decreased by 174- to over 38,000-fold compared to mice infected with WT. Taken together, these data demonstrate that most of the *L. monocytogenes* genes identified by Tn-seq as important for colonization of NHP gallbladders *ex vivo* were also required for infection of murine gallbladders *in vivo*.

In addition to the gallbladders, bacterial burdens were measured from infected livers, mesenteric lymph nodes (MLN), and spleens. After ingestion, *L. monocytogenes* in the GI tract disseminates via the portal vein to the liver and via the lymphatics through the MLN and to the spleen [5]. Bacterial burdens in the livers were similar between the mutants and WT at 1 dpi (S3A Fig), further demonstrating that these genes are not required for dissemination via the portal vein to the liver. At 4 dpi, all mutants (except *ΔccpA*) displayed significantly decreased bacterial burdens in the livers compared to WT (Fig 4E). Most mutants colonized the MLN and displayed similar bacterial burdens as WT at both 1 and 4 dpi. In contrast, *ΔptsI*, *ΔpurB*, and *ΔatpB* were significantly attenuated in the MLN compared to WT (S3C and S3D Figs). In the spleens, only *ΔptsI, ΔpurB, Δmpt,* and *clpX::Tn* were significantly attenuated at 4 dpi compared to WT (Fig 4F). The primary source of *L. monocytogenes* excreted in the feces is derived from the population in the gallbladder; thus, bacterial burdens in the feces were also measured [7]. Bacterial burdens in the feces were similar between most mutants and WT at 1 dpi, whereas the majority of mutants exhibited significantly decreased bacterial loads compared to WT at 4 dpi (S3E and S3F Figs). These data collectively demonstrate that the genes identified in our *ex vivo* screen contribute to infection of multiple organs, including the gallbladder, following oral infection of mice.

## DISCUSSION

In this study we sought to identify *L. monocytogenes* genes important for infection of the mammalian gallbladder. To this end, we developed an *ex vivo* bacterial colonization model of NHP gallbladders and performed Tn-seq to determine the genes necessary for growth and survival in this environment. This unbiased global genetic approach identified mutants in 43 genes that were significantly depleted after growth in the gallbladder condition, including some genes known to be important for virulence and others not previously studied in the context of infection. Several mutants identified by Tn-seq had growth defects in rich medium and most were predictably attenuated for growth in NHP bile *in vitro*. Many mutants also had defects in the intracellular lifecycle, including cytosolic growth and cell-to-cell spread, with the notable exception of the PTS permeases. A murine model of oral *L. monocytogenes* infection revealed that nearly all identified genes are required for full virulence. Together, these data identified genes that are important for *L. monocytogenes* infection of a mammal and, interestingly, not all are required for intracellular replication or intercellular spread.

Animal models have shed light on the importance of gallbladder colonization during *L. monocytogenes* pathogenesis. Murine models of infection demonstrated that *L. monocytogenes* replicates extracellularly in the gallbladder lumen to high bacterial densities and that this population becomes the primary bacterial reservoir and source of fecally shed *L. monocytogenes* [4,7,10]. Further, they revealed the presence of an uncharacterized severe within-host bottleneck in which the founding population of the gallbladder is limited to approximately 3 bacteria [7]. A study using *ex vivo* porcine gallbladders found that *L. monocytogenes* readily grows in both the whole organ and extracted bile [17]. Given the limitations of using murine or porcine infection models for examining bacterial gallbladder colonization, we established a new infection model with a less restrictive bottleneck that is also amenable to Tn-seq analysis. While the present study focused on *L. monocytogenes* luminal growth within the organs, the *ex vivo* NHP gallbladder model could be utilized to measure mucosal and epithelial colonization of a variety of gallbladder-tropic pathogens.

Our Tn-seq screen identified mutants in dozens of genes that were significantly depleted after incubation in the NHP gallbladders. These genes include those involved in redox regulation (*trxA, yjbH, rex*), cell wall modifications (*walK*), and protein stability (*clpX*), consistent with the known antimicrobial activities of bile which result in cell envelope stress and protein damage [9]. Purine biosynthesis was previously identified as important for growth in porcine bile and here, we identified *purB* as essential for replication in the gallbladder [17]. Interestingly, the screen did not identify *mdrT*, *bsh*, *bilE*, or *sigB* as required for growth or survival in the gallbladder, despite previous reports that they are all essential for *L. monocytogenes* replication in bile [16,29,30]. MdrT is a multidrug resistance transporter originally identified to secrete c-di-AMP [31,32] and subsequently suggested to be an efflux pump for cholic acid, a component of bile [33]. The *bsh* gene encoding <u>b</u>ile <u>s</u>alt <u>h</u>ydrolase was originally described as required for survival in bile *in vitro* and for intestinal persistence in a guinea pig model of infection [29]. The <u>bil</u>e <u>e</u>xclusion system encoded by *bilE* was proposed to be a transporter that protected *L. monocytogenes* from toxicity induced by 30% reconstituted bovine bile [30]. SigB is a stress response alternative sigma factor that positively regulates both *bsh* and *bilE* [16,30]. It is now appreciated that *bsh, bilE*, and *sigB* confer resistance to acidified bile acids, as may be found in the small intestine, but are not necessary to detoxify bile at neutral pH, as would be found in the gallbladder lumen [17,34]. Furthermore, BilE was recently renamed EgtU when it was conclusively demonstrated that it specifically binds and transports the low molecular weight thiol ergothioneine [35].

Some genes we identified by Tn-seq had been previously described as necessary for virulence *in vivo,* while many others had not been studied in the context of infection. The *purB, trxA, yjbH, clpX*, and *rex* genes are known to be required for full virulence, though some of the studies used different mouse strains and different inoculation methods [36–39]. Moreover, *rex* is one of the few *L. monocytogenes* genes specifically required for replication in the murine gallbladder [34]. Conversely, we identified genes encoding all 8 structural components of the F-type ATP synthase, which was not previously examined *in vivo* [25]. We also identified operons encoding two PTS EII complexes (*mpt, mpo*), as well as genes encoding their transcriptional (*sigL, manR*) and post-transcriptional regulators (*ptsH, ptsI*). Mpt and Mpo were previously designated as dispensable for virulence base on tissue culture assays, although were never tested *in vivo* [23].

Most mutants under investigation were deficient for growth in NHP bile *in vitro*, which was unsurprising given the conditions under which the Tn-seq screen was performed. The most attenuated strains after growth in bile were *ΔptsI*, *ΔtrxA*, and *ΔpurB*, which also displayed growth defects in rich medium. Additionally, the *ΔatpB* strain was severely attenuated in BHI, which was expected based on the published requirement for the F-type ATP synthase for anaerobic replication [25]. In the aerobic condition, the cultures were incubated statically and we hypothesize that the lack of aeration led to *ΔatpB* growing more slowly than WT in BHI. Surprisingly, *ΔatpB* did replicate in NHP bile under anaerobic conditions. Future research will investigate the role of *atpB* in *L. monocytogenes* growth in bile and *in vivo*. It has been hypothesized that the F-type ATP synthase is required during anaerobic to combat acid stress and generate a proton motive force, rather than for ATP synthesis [25,40]. It remains unclear, however, if these mechanisms contribute to the role of the F-type ATP synthase during infection.

Tissue culture models of infection have historically been reliable indicators of *L. monocytogenes* pathogenesis *in vivo*, although the correlation was not as strong in this study. Several mutants displayed defects in a plaque assay, which measures both intracellular growth and intercellular spread over three days. Specifically, Δ*ccpA, ΔtrxA, ΔpurB*, and *clpX::Tn* formed significantly smaller plaques than WT. Despite this, Δ*ccpA* was surprisingly fully virulent in a murine model of listeriosis. Furthermore, *mpt* and *mpo* were completely dispensable for intracellular growth and intercellular spread in tissue culture, although they were required for infection of mice. Importantly, studies solely using tissue culture models of infection would not have identified these operons as important for pathogenesis *in vivo*.

To assess the validity of the *ex vivo* NHP gallbladder model, we evaluated the roles of genes identified by Tn-seq in an oral murine model of listeriosis. Nearly all mutants tested were significantly attenuated in the gallbladders and livers of infected mice 4 dpi. The notable exception was the strain lacking *ccpA,* the <u>c</u>atabolite <u>c</u>ontrol <u>p</u>rotein <u>A</u> that represses transcription of metabolic genes based on the phosphorylation state of HPr and the overall nutrient status of the cell. In the Δ*ccpA* mutant, approximately 100 genes are de-repressed [41], resulting in attenuated *in vitro* growth in rich medium, bile, and BMDMs, yet exhibiting no virulence defect in mice after oral infection. Interestingly, not all strains were attenuated in the spleens and MLNs of infected mice, suggesting that factors required for colonizing the liver and gallbladder are distinct from those needed to colonize other peripheral organs. For example, Δ*trxA* was attenuated approximately 29,000-fold in the gallbladder, but colonized the spleen and MLN at levels similar to WT. Relatedly, it was recently reported that *L. monocytogenes* folate metabolism is specifically required in the livers but not the spleens of infected mice [22,42,43]. Similarly to the gallbladders, most mutants were significantly attenuated in the feces at 4 dpi, supporting the notion that the gallbladder is the main source of fecally excreted bacteria [7]. The two mutants not attenuated in the feces were *ΔccpA* and Δ*purB.* While Δ*ccpA* was fully virulent in mice, the Δ*purB* mutant was severely attenuated *in vitro* and in all murine organs after infection. These results indicate that purine biosynthesis is required for intracellular infection and virulence, but not for survival in the GI tract, consistent with previous reports [36,39]. Taken together, oral infections of mice revealed that the genes identified by Tn-seq in the NHP gallbladder have broader roles in disease pathogenesis than simply conferring resistance to bile stress.

Taken together, our Tn-seq approach revealed several novel insights into *L. monocytogenes* carbon metabolism during infection. It is well-accepted that the primary carbon sources consumed by *L. monocytogenes* in the cytosol are host-derived glycerol and hexose-phosphates and thus, the permeases that import these sugars are required for intracellular replication and virulence [44–46]. Conversely, *L. monocytogenes* encodes 84 genes that assemble into 29 complete PTSs, which were previously thought to be dispensable for virulence [23]. Indeed, the main glucose and mannose PTS EII proteins, encoded by the *mpt* and *mpo* operons, are not required for intracellular growth or intercellular spread [23,47]. However, we found that strains lacking *mpt* or *mpo* are significantly attenuated in a murine model of listeriosis. These results suggest that glucose and mannose are important nutrients for *L. monocytogenes* replicating in extracellular sites *in vivo*, such as the gallbladder and, to a lesser extent, the liver. Recent studies established that significant populations of *L. monocytogenes* are extracellular in the liver, spleen, and MLN [6,48], although the bacterial requirements for surviving extracellularly and the role that extracellular bacteria play in pathogenesis remain unclear. Moreover, all PTSs are activated by a phosphorelay between Enzyme I (encoded by *ptsI*) and HPr (encoded by *ptsH*). Thus, the Δ*ptsI* mutant, which functionally lacks all 29 PTSs, is deficient for intracellular replication and dramatically attenuated *in vivo.* This suggests that PTS-dependent carbohydrates are important nutrients in the host cytosol. Ongoing studies are aimed at characterizing the additional PTSs that are required for full virulence.

## METHODS

### Ethics Statement

This study was carried out in strict accordance with the recommendations in the Guide for the Care and Use of Laboratory Animals of the National Institutes of Health. All protocols were reviewed and approved by the Animal Care and Use Committee at the University of Washington (Protocol 4410– 01).

### Bacterial strains and conditions

The bacterial strains used in this study are listed in S2 Table. *L. monocytogenes* was cultured in brain heart infusion (BHI) and *E. coli* was cultured in Luria-Bertani (LB) broth at 37°C, with shaking (220 rpm), unless otherwise specified. Antibiotics (purchased from Sigma Aldrich) were used at the following concentrations: streptomycin, 200 μg/mL; chloramphenicol, 10 μg/mL (*E. coli*) and 7.5 μg/mL (*L. monocytogenes*); and carbenicillin, 100 μg/mL. *L. monocytogenes* mutants were derived from wild type strain 10403S [49,50]. Plasmids were introduced to *E. coli* via chemical competence and heat shock and introduced into *L. monocytogenes* via trans-conjugation from *E. coli* SM10 [51].

### Vector construction and cloning

To construct deletion mutants in *L. monocytogenes*, ∼700 bp regions up- and downstream of the gene of interest were PCR amplified using *L. monocytogenes* 10403S genomic DNA as a template. PCR products were digested and ligated into pLIM (gift from Arne Rietsche, Case Western). pLIM plasmids were then transformed into *E. coli* and sequences confirmed via Sanger sequencing (Azenta). Plasmids harboring mutant alleles were then introduced into *L. monocytogenes* via trans-conjugation and integrated into the chromosome as previously described [52,53].

Complemented strains of *L. monocytogenes* were generated using the pPL2 integration plasmid [54]. Genes were PCR amplified with their respective native promoters using *L. monocytogenes* 10403S genomic DNA as a template, and sequences were confirmed by Sanger sequencing. The constructed pPL2 plasmids were then introduced into *L. monocytogenes* by trans-conjugation and integration into the *L. monocytogenes* chromosome was confirmed by antibiotic resistance.

### Growth curves in NHP bile

NHP bile aliquots were plated to evaluate sterility and stored at -80°C. Before an experiment, aliquots were thawed overnight at 4°C and warmed to room temperature immediately before the inoculation into a 96-well plate. Overnight *L. monocytogenes* cultures were washed twice, resuspended in PBS, and 10^5^ CFU were inoculated into either NHP bile or BHI, in a total volume of 100 µL per well. Bacterial growth was measured by collecting samples of the cultures, serially diluting in PBS, and plating for CFU. For experiments performed anaerobically, NHP bile aliquots were thawed overnight in GasPak EZ Anaerobe gas-generating pouches (Becton Dickinson), and BHI and the 96-well plate were degassed overnight in a closed-system anaerobic chamber (Don Whitley Scientific A35 anaerobic work station). After washing and resuspending aerobically-grown overnight *L. monocytogenes* cultures in PBS, the *L. monocytogenes* suspensions and bile aliquots were transferred into the anaerobic chamber and the plate was inoculated and incubated within the chamber. Bacterial growth was measured by collecting samples of the cultures, serially diluting in PBS, and plating for CFU.

### *L. monocytogenes* transposon library in NHP gallbladders

The *L. monocytogenes* transposon library [22] was inoculated directly from the -80°C stock into BHI broth and incubated at 37°C for 2 hours, with shaking. The library was then washed twice and resuspended in PBS to a density of 10^8^ CFU per 2 kg of NHP body weight. The inoculum size was determined to maintain 1,000-fold coverage of the library. Gallbladders were injected via syringe with 100 µL of inoculum, the injection site was sealed with liquid bandage (3M), and incubated in a 15 cm petri dish at 37°C and 5% CO_2_. After 30 minutes, 200 µL of bile was removed from the gallbladder via syringe, serially diluted, and plated to enumerate CFU. 6 hours post-injection, bile was extracted from the organ via syringe and the remaining luminal contents collected via cell scraper after resection. The gallbladder contents were diluted into 50 mL BHI broth and incubated at 37°C for 2 hours, with shaking. The cultures were pelleted, washed twice with PBS, and stored at -80°C.

### Tn-seq library preparation, sequencing, and analysis

Genomic DNA was extracted using a Quick-DNA Fungal/Bacterial MiniPrep Kit (Zymo Research). DNA was diluted to 3 µg/130 µL in microTUBES (Covaris) and sheared in duplicate on a Covaris LE220 Focused-Ultrasonicator using the following settings: duty cycle 10%; peak intensity 450; cycles per burst 100; duration 100 sec. Sheared DNA was then end-repaired with NEBNext End Repair (NEB), and purified with Ampure SPRIselect beads (Beckman Coulter). Poly-C tails were added to 1 µg of end-repaired DNA with Terminal Transferase (Promega), then purified with Ampure SPRIselect beads. Transposon junctions were PCR amplified with primers olj376 and pJZ_RND1 (Sx Table) using 500 ng DNA and KAPA HiFi Hotstart Mix (Kapa Biosystems). PCR reactions were stopped once the inflection point of amplification was reached (6-14 cycles), and amplified transposon junctions were purified with Ampure SPRIselect beads. Barcoded adaptors were added using KAPA HiFi Hotstart Mix, and primers pJZ_RND2 and one TdT_Index per sample. DNA was purified and size-selected with Ampure SPRIselect beads for 250-450 bp fragments. Samples were pooled and sequenced as single end 50 bp reads on a NextSeq MO150 sequencer with a 7% PhiX spike in and primer pJZTnSq_SeqPrimer.

Trimmed reads were mapped to the *L. monocytogenes* 10403S NC_17544 reference genome in PATRIC (now https://www.bv-brc.org/) and assessed for essentiality using TRANSIT software [55–57]. Genes were considered required for survival or growth in *ex vivo* NHP gallbladders if they met the following criteria: 5 or more insertion sites in the input libraries, a *p* value less than 0.05, and a 1.5-fold or greater depletion after incubation in the gallbladder.

### Murine cells

L2 fibroblasts were incubated at 37°C in 5% CO2 in Dulbecco’s modified Eagle’s medium (DMEM) with 10% heat-inactivated fetal bovine serum (FBS) (Cytiva) and supplemented with sodium pyruvate (1 mM) and L-glutamine (1 mM) (L2 Medium). For passaging, cells were maintained in Pen-Strep (100 U/ml) but were plated in antibiotic-free media for infections.

Bone marrow-derived macrophages (BMDMs) were routinely incubated in DMEM supplemented with 20% heat-inactivated FBS, 1 mM sodium pyruvate, 1 mM L-glutamine, 10% supernatant from M-CSF-producing 3T3 cells, and 55 μM β-mercaptoethanol (BMDM medium). BMDMs were isolated as previously described [58]. Briefly, femurs and tibias from C57BL/6 mice were crushed with a mortar and pestle in 20 mL BMDM medium and strained through 70-μm cell strainers. Cells were plated in 150-mm untreated culture dishes, supplemented with fresh BMDM medium at day 3, and then harvested by resuspending cells in cold PBS at day 7. BMDMs were aliquoted in 80% BMDM medium, 10% FBS, and 10% DMSO and stored in liquid nitrogen.

### Intracellular growth curves

BMDMs were plated in TC-treated 24-well plates at a density of 6 x 10^5^ cells per well in BMDM medium. *L. monocytogenes* cultures were grown overnight at 30°C, stationary. The next day, *L. monocytogenes* cultures were washed twice, resuspended in PBS, and added to BMDMs at an MOI = 0.1. After 30 minutes, cells were washed twice with PBS and BMDM medium containing gentamicin (50 µg/mL) was added to kill extracellular bacteria. At various time points post-infection, cells were washed twice with PBS and lysed in 250 μL cold 0.1% Triton-X in PBS. Lysates were then serially diluted and plated to enumerate intracellular CFU.

### Plaque assays

Plaque assays were performed as previously described [38,59]. In brief, 1.2 x 10^6^ L2 fibroblasts were plated in tissue-culture treated 6-well plates overnight in L2 medium. *L. monocytogenes* cultures were grown overnight at 30°C stationary. The next day, *L. monocytogenes* cultures were diluted 1:10 in PBS and 5 µL of diluted bacteria was added to cell monolayers. After 1 hour of infection, monolayers were washed twice with PBS, then overlaid with 3 mL of molten agarose solution (1:1 mixture of 2X DMEM and 1.4% SuperPure Agarose (U.S. Biotech Sources, LLC), containing 10 µg/mL gentamicin). After 3 days of incubation, 2 mL of molten agarose solution containing Neutral Red was added to wells to visualize plaques. After 12-24 hours, plates were scanned, plaque areas quantified using ImageJ software [60] and normalized to WT.

### Oral murine infections

Female BALB/c mice were purchased from The Jackson Laboratory at 5 weeks of age and used in experiments when they were 6–7 weeks old. Infections were performed as previously described [4,34]. Streptomycin (5 mg/mL) was added to drinking water 48 hours prior to infection and food and water were removed 16 hours before infection. *L. monocytogenes* cultures were grown overnight at 30°C, stationary. Overnight cultures were diluted 1:10 in 5 mL fresh BHI and incubated at 37°C for 2 hours, with shaking. Bacteria were then washed twice and diluted in PBS. Mice were fed 10^8^ bacteria in 20 µL of PBS and food and water were returned immediately after infection. Inocula were serially diluted and plated. Body weights were recorded daily and mice were humanely euthanized 1 and 4 days post-infection for tissue collection. Tissues were homogenized in the following volumes of 0.1% Igepal CA-630 (Sigma): MLN, 3 mL; cecum (contents removed and tissues rinsed with PBS), 4 mL; liver, 5 mL; spleen, 3 mL. Feces were homogenized in 1 mL of 0.1% Igepal with a sterile stick, and gallbladders were ruptured and crushed in 500 µL of 0.1% Igepal with a sterile stick. All samples were serially diluted in PBS and plated to enumerate CFU.

## ACKNOWLEDGEMENTS

We would like to thank all members of the Reniere and Woodward Labs at the University of Washington (UW) for valuable feedback and discussions; the Mougous Lab at UW for equipment; and Christopher English at the Washington National Primate Research Center for coordinating NHP experiments.

## SUPPORTING INFORMATION LEGENDS

**S1 Figure. Growth curves of *L. monocytogenes* in BHI or bile *in vitro*.** BHI or NHP bile was inoculated with *L. monocytogenes*, incubated statically in an aerobic incubator (A,B) or in an anaerobic chamber (C,D), and CFU were enumerated, as in Figure 2. Data are the means and SEMs of at least three independent experiments. Statistics not shown for clarity.

**S2 Figure. Intracellular replication of complemented strains.** Intracellular growth of complemented mutants in BMDMs, normalized to CFU at 30 minutes for each strain. Data are the means and SEMs of two independent experiments.

**S3 Figure. Murine model of oral listeriosis.** Mice were orally infected with 10^8^ of each *L. monocytogenes* strain and CFU from tissues or feces were enumerated 1 (A, B, C, E) or 4 (D, F) days post-infection. Each data point represents a single mouse (n=15 for WT, n=4-10 for mutants), horizontal solid lines represent geometric means, and the dotted lines represent the limit of detection. The data are combined from 4 independent experiments. * *p* < 0.05, ** *p* < 0.01, *** *p* < 0.001, as determined by Dunnett’s multiple comparisons test compared to WT in each condition.

**S1 Table. Tn-seq results**

**S2 Table. Strains used in this study**

**S3 Table. Primers used in Tn-seq library preparation**

